# β-Adrenergic Stimulation Synchronizes a Broad Spectrum of Action Potential Firing Rates of Cardiac Pacemaker Cells towards a Higher Population Average

**DOI:** 10.1101/2021.07.06.451302

**Authors:** Mary S. Kim, Oliver Monfredi, Larissa A. Maltseva, Edward G. Lakatta, Victor A. Maltsev

## Abstract

The heartbeat is initiated by pacemaker cells residing in the sinoatrial node (SAN). SAN cells generate spontaneous action potentials (APs), i.e. normal automaticity. The sympathetic nervous system increases heart rate commensurate with cardiac output demand *via* stimulation of SAN β-adrenergic receptors (βAR). While SAN cells reportedly represent a highly heterogeneous cell population, the current dogma is that in response to βAR stimulation all cells increase their spontaneous AP firing rate in a similar fashion. The aim of the present study was to investigate cell-to-cell variability in the responses of a large population of SAN cells. We measured βAR responses among 166 single SAN cells isolated from 33 guinea pig hearts. In contrast to the current dogma, the SAN cell responses to βAR stimulation substantially varied. In each cell, changes in AP cycle length highly correlated (R^2^=0.97) with the AP cycle length before βAR stimulation. While, as expected, on average the cells increased their pacemaker rate, greater responses were observed in cells with slower basal rates, and vice versa, cells with higher basal rates showed smaller responses, no responses, or even decreased their rate. Thus, βAR stimulation *synchronizes* the operation of the SAN cell population towards a higher average rate, rather than uniformly shifting the rate in each cell, creating a new paradigm of βAR-driven fight-or-flight response among individual pacemaker cells.

## 1. Introduction

Enzymatically isolated single sinoatrial node (SAN) cells have been extensively utilized as a model in studies of cellular mechanisms of cardiac pacemaker rate regulation [1]. These studies have advanced our understanding of how the heart responds to autonomic receptor stimulation: β-adrenergic receptor (βAR) stimulation increases spontaneous action potential (AP) firing rate in SAN, while cholinergic receptor stimulation decreases the rate. Further, it is generally believed that βAR stimulation shifts AP firing rate in a similar fashion in all SAN cells *via* a small increase in the slope of diastolic depolarization [2]. This idea is based on electrophysiological measurements performed in a highly selected, relatively small population of cells. These selected cells, previously believed to be “true” pacemaker cells, contract spontaneously, and beat frequently and rhythmically after enzymatic isolation, mirroring the behavior of the whole SA node. However, only 10-30% of isolated cells behaved in this way in the original paper describing SAN cell isolation [3]. The isolation procedure has been refined and improved over time, and while the yield of spontaneously and rhythmically contracting cells has increased, it has never approached 100%. In addition to cells firing frequent and rhythmic spontaneous APs, many isolated cells fire APs at slower, sometimes irregular rates, while others remain silent and do not beat at all despite a normal appearance.

Traditional research focusing only on the frequently beating “true” SAN cells has been justified on the basis of the operational paradigm that SAN tissue is driven by a leading pacemaker cell (or cluster) that dictates excitation rate and rhythm to all other subservient cells [4,5]. Coordinated firing of individual cells within SAN tissue has also been approached along the lines of mutual entrainment of individual cells towards a single rate within a network of loosely coupled oscillators (dubbed the ‘democratic process’) [6,7]. As a result, historic SAN cell studies have deliberately selected only those cells that are spontaneously beating after isolation, with rates and rhythms similar to that of intact SAN tissue. All other cells remaining after enzymatic isolation were typically ignored. This includes non-beating, irregular or infrequently beating cells, have been typically rejected for study on the basis that they were “damaged” by the enzymatic isolation procedure, or that they were subsidiary or not “true” pacemaker cells.

This “one rate/rhythm” paradigm of SAN cell operation, and the related criteria of pacemaker cell vitality and authenticity, has recently been challenged. Using a novel high-resolution imaging technique, Bychkov et al. [8] recorded Ca signals in individual cells over the entire intact mouse SAN. They found that synchronized cardiac impulses emerge from heterogeneous local Ca signals within and among cells of SAN tissue. The patterns seen resembled complex processes of impulse generation within clusters of neurons in neuronal networks. Many cells within the HCN4-positive network (where pacemaker impulses first emerge) in fact did not generate APs at all, or generated APs with rates and rhythms different from those exiting the SAN to capture the atria. Another recent study [9] in knock-in mice expressing cAMP-insensitive HCN4 channels showed that tonic and mutual interaction (so-called ‘tonic entrainment’) between firing and non-firing cells slows down SAN pacemaking. cAMP increased the proportion of firing cells in this work, and in doing so increased the rate of SAN automaticity, and increased resistance to parasympathetic stimulation.

We have recently examined non-firing, silent cells - “dormant cells” - isolated from guinea pig and human hearts [10–12]. Many dormant cells can be reversibly “awakened” to fire normal, spontaneous rhythmic APs by βAR stimulation. The transition to spontaneous AP firing occurs via the emergence and coupling of rhythmic intracellular Ca oscillations (known as the Ca clock [13,14]). These rhythmic local Ca releases (LCRs) interact with the membrane oscillator (known as the membrane clock) to drive the diastolic depolarization and trigger high-amplitude APs. During the transition to AP firing, both clocks’ functions undergo specific changes including: (i) increased cAMP-mediated If activation; (ii) increased cAMP-mediated phosphorylation of phospholamban; this increases ICaL density and accelerates Ca pumping; and (iii) increased spatiotemporal LCR synchronization; this yields a larger diastolic LCR ensemble signal and an earlier increase in diastolic Na/Ca exchanger current [12]. All these changes result in effective AP ignition required for Ca and membrane clock coupling during diastolic depolarization [15].

While cAMP-dependent phosphorylation is a key regulator of clock coupling [16], how different degrees of phosphorylation translate into clock coupling and cell behavior (e.g. dormancy, rhythmicity or dysrhythmic firing, along with effects on autonomic receptor modulation, etc.) will depend on the expression level of clock proteins in individual cells. This cannot be assumed to be the same from cell to cell. Indeed, immunocytochemical labeling of the L-type Ca channels, Na/Ca exchanger, Ca release channels (RyR2), and Ca pump (SERCA2) varies widely in SAN cells [17]. SAN cells also exhibit a substantial degree of cell-to-cell variability in functional expression of ICaL, If, and IK [18,19]. For example, cell-to-cell variations in ICaL or If densities can be as much as an order of magnitude. LCR characteristics also vary substantially between cells, with the LCR period highly correlating with the AP cycle length [20].We know from many years of observation that enzymatically isolated SAN cells have a wide spectrum of basal beating rates (from high beating rates all the way to dormancy), and that this could be due to natural diversity rather than damage during isolation procedure.

The aim of the current study was, for the first time, to document the range of responses of all spontaneously beating isolated SAN cells to βAR stimulation, regardless of their basal beating rate. We hypothesized that there would be a wide variety of responses to βAR stimulation in heterogenous isolated SAN cells, and that this would be related to the basal beating rate pre-βAR stimulation.

We demonstrate that SAN cell responses to βAR stimulation do indeed differ fundamentally from the dogma that all cells increase their rate in a similar fashion. Most cells did increase their rate, but some were insensitive to stimulation, and others even paradoxically decreased their rates. Overall, during βAR stimulation, the spectrum of rates among individual cells substantially narrowed, becoming synchronized around a higher average.

## 2. Materials and Methods

### 2.1. Single cell preparation

SAN cells were isolated from 33 male guinea pigs in accordance with NIH guidelines for the care and use of animals, protocol 034-LCS-2019 [10]. Hartley guinea pigs (Charles River Laboratories, USA) weighing 500-650 g were anesthetized with sodium pentobarbital (50–90 mg/kg). The heart was removed quickly and placed in solution containing (in mM): 130 NaCl, 24 NaHCO_3_, 1.2 NaH_2_PO_4_, 1.0 MgCl_2_, 1.8 CaCl_2_, 4.0 KCl, 5.6 glucose equilibrated with 95% O_2_ / 5% CO_2_ (pH 7.4 at 35°C). The SA node region was cut into small strips (~1.0 mm wide) perpendicular to the crista terminalis and excised. The final SA node preparation consisted of SA node strips attached to the small portion of crista terminalis. The SA node preparation was washed twice in Ca-free solution containing (in mM): 140 NaCl, 5.4 KCl, 0.5 MgCl_2_, 0.33 NaH_2_PO_4_, 5 HEPES, 5.5 glucose, (pH=6.9) and incubated on a shaker at 35°C for 30 min in the same solution with the addition of elastase type IV (0.6 mg/ml of 5.7 units per mg; Sigma, Chemical Co.), collagenase type 2 (0.8 mg/ml of 250 units per mg; Worthington, NJ, USA), protease XIV (0.12 mg/ml of ≥3.5 units per mg; Sigma, Chemical Co.), and 0.1% bovine serum albumin (Sigma, Chemical Co.).

The SAN preparation was washed in modified Kraftbruhe (KB) solution, containing (in mM): 70 potassium glutamate, 30 KCl, 10 KH_2_PO_4_, 1 MgCl_2_, 20 taurine, 10 glucose, 0.3 EGTA, and 10 HEPES (titrated to pH 7.4 with KOH), and kept at 4°C for 1h in KB solution containing 50 mg/ml polyvinylpyrrolidone (Sigma, Chemical Co.). Finally, cells were dispersed from the SA node preparation by gentle pipetting in the KB solution and stored at 4°C for subsequent use in our experiments for 8 hours.

The cells measured in this study most likely correspond to sinus node pacemaker cells rather than atrial cells because of the following:

- We accurately dissected the SA node to avoid (or minimize) contamination with atrial myocytes. The location of guinea pig SA node in relation to the whole heart during our isolation procedure is shown in Figure S1.
- We measured SA node cells that had a classical spindle-shaped morphology (Figure S2)
- We deliberately avoided cells with typical atrial cell morphology (i.e. blocky and rectangular in shape).
- We described electrophysiological properties of these cells in our recent study [12]. Cells used in the present study have similar properties because they were from the same isolations (from the same guinea pig hearts). The maximum diastolic potential was on average −58.5 mV. This is clearly different from SA node-residing atrial cells that have a stable resting membrane potential between −70 and −80 mV [21].

### 2.2. Two-dimensional (2D) Ca imaging of single cells

Ca dynamics within isolated single SAN cells were measured by 2D imaging of fluorescence emitted by the Ca indicator Fluo-4 (Invitrogen) using a Hamamatsu C9100-12 CCD camera, with an 8.192 mm square sensor of 512 × 512 pixels resolution), as previously described [22]. The sampling rate of 100 frames/sec was chosen for our imaging because it satisfactorily reports both the AP cycle length and spatiotemporal resolution of LCRs [22]. The camera was mounted on a Zeiss Axiovert 100 inverted microscope (Carl Zeiss, Inc., Germany) with x63 oil immersion lens and a fluorescence excitation light source CoolLED pE-300-W (CoolLED Ltd. Andover, UK). Fluo-4 fluorescence excitation (blue light, 470/40 nm) and emission light collection (green light, 525/50 nm) were performed using the Zeiss filter set 38 HE. Cells were loaded with 5 mM Fluo-4AM (Sigma-Aldrich, USA) for 20 minutes at room temperature. The physiological (bathing) solution contained (in mM): 140 NaCl; 5.4 KCl; 2 MgCl_2_; 5 HEPES; 1.8 CaCl_2_; pH 7.3 (adjusted with NaOH). Data acquisition was performed using SimplePCI (Hamamatsu Corporation, Japan) at physiological temperature of 35°C ± 0.1°C.

### 2.3. Detection and analysis of the whole ensemble of AP-induced cytosolic Ca transients and LCRs in 2D Ca imaging

AP-induced Ca transients were observed as sharp, whole-cell wide (i.e. within the entire cell perimeter) transient rises of Fluo-4 fluorescence. We considered the time periods between these high amplitude signals as a satisfactory measure of AP cycle length (further referred as cycle length) as we previously reported in simultaneous recordings of Ca and membrane potential [23]. The time series of whole-cell Fluo-4 fluorescence (within the cell perimeter) were measured from 2D video recordings using HCImageLive image analysis software (v.3, x64, Hamamatsu Co., Japan). Then fluorescence peaks (i.e. AP-induced Ca transients) and their respective cycle lengths were determined by our custom-made program (Victor Maltsev).

LCRs were observed as local Ca signals scattered within cell perimeter. They were detected and analyzed by program “XYT Event Detector” [24] (its C++ code is freely available on NIH website https://www.nia.nih.gov/research/labs/xyt-event-detector)

In short, the program detects LCR birth/death events by a differential, frame-to-frame sensitivity algorithm applied to each pixel (cell location) in a series of images generated by the camera. An LCR is detected when its signal changes sufficiently quickly within a sufficiently large area. The LCR “dies” when its amplitude decays substantially or when it merges into a rising AP-induced Ca transient. LCRs were isolated from noise by applying a series of spatial filters that set the minimum pixel size and the brightness threshold for LCRs. Cells with mechanical contractile movements were affixed by tracking points along the midline of the moving cell by using a computer program “SANC Analysis” described in [24]. The algorithm uses these points as a coordinate system for affine transform, producing a transformed image series that are stationary. The program is available as a free ImageJ plugin on the website http://scepticalphysiologist.com/code/code.html created and maintained by Dr. Sean Parsons.

### 2.4. Evaluation of βAR stimulation effect

βARs were stimulated by 1 μM isoproterenol in SAN cells during continuous perfusion of the entire bath with physiological solution. In each cell Ca signals were recorded two times: before βAR stimulation (designated as control or basal state) and during stimulation approximately 5 minutes after isoproterenol application onset. To avoid photodamage each recording was limited to 30 seconds. The respective cycle lengths were measured from each recording (as described above) and each measured cell was represented by two numbers: average cycle lengths in control and in the presence of βAR stimulation. We also evaluated effect of βAR stimulation on LCR characteristics, namely: LCR period, time from AP-induced Ca transient peak to the onset of an LCR (ms); LCR duration, time from LCR onset to its “death” or merging to the transient (ms); LCR area, the cell area that was covered by a given LCR during its life span including expansion and propagation (μm^2^) (reflecting parameter “LCR spatial size” reported in confocal line-scan images [20]); maximum LCR size, the area of the largest LCR in each cycle (μm^2^).

### 2.5. Statistics

Data are presented as mean ± SEM. The statistical significance of the effects of βAR stimulation on cycle length and LCR characteristics was evaluated by paired t-test (n=166, in each cell before and during stimulation) using Data Analysis Add-In of Microsoft Excel program, version 2106.

## 3. Results

### 3.1. Cycle length changes

On average, SAN cells significantly decreased their spontaneous cycle length in response to βAR stimulation. Statistical evaluation of the responses is provided in Table 1.

**Table 1.** Effect of βAR stimulation on cycle length measured in a population of 166 SAN cells that generated spontaneous AP-induced Ca transients; SEM, standard error of mean; SD, standard deviation, CV%= 100* SD/Mean, coefficient of variation.

| Parameter | Before stimulation | βAR stimulation |
| --- | --- | --- |
| Mean cycle length, ms | 921.8<sup>1</sup> | 507.3<sup>1</sup> |
| SEM, ms | 66.5 | 13.1 |
| SD, ms | 856.4 | 169.1 |
| CV% | 92.9 | 33.3 |

<sup>1</sup> Statistically significant with $P < 0.001$ *via* paired t-test, two tail.

Responses varied substantially from cell to cell and can be categorized into three types: (i) Most cells, 121 of 166 (72.9%), showed the “classical” response of cycle length reduction, i.e. their rate increased; (ii) a smaller fraction of cells, 12 of 166 (7.2%) showed almost no change in cycle length (cycle length remained within 3% of baseline); (iii) the remainder, 33 of 166 (19.9%) showed the “unexpected” or “paradoxical” response of cycle length increase (AP firing rates decrease). Examples of each category are shown in Figure 1 and respective Videos S1-S6 (each response category is represented by two video recordings: before βAR stimulation and during stimulation. Another way to categorize cell responses is to exclude the “no change” category and formally divide all cells based on the sign of the cycle length change. In this case we report 38 cells of 166 (i.e. 22.9%) that responded with cycle length increase (Table S1). Anyway, the pool of cells with the unusual response is notable.

**Figure 1.**
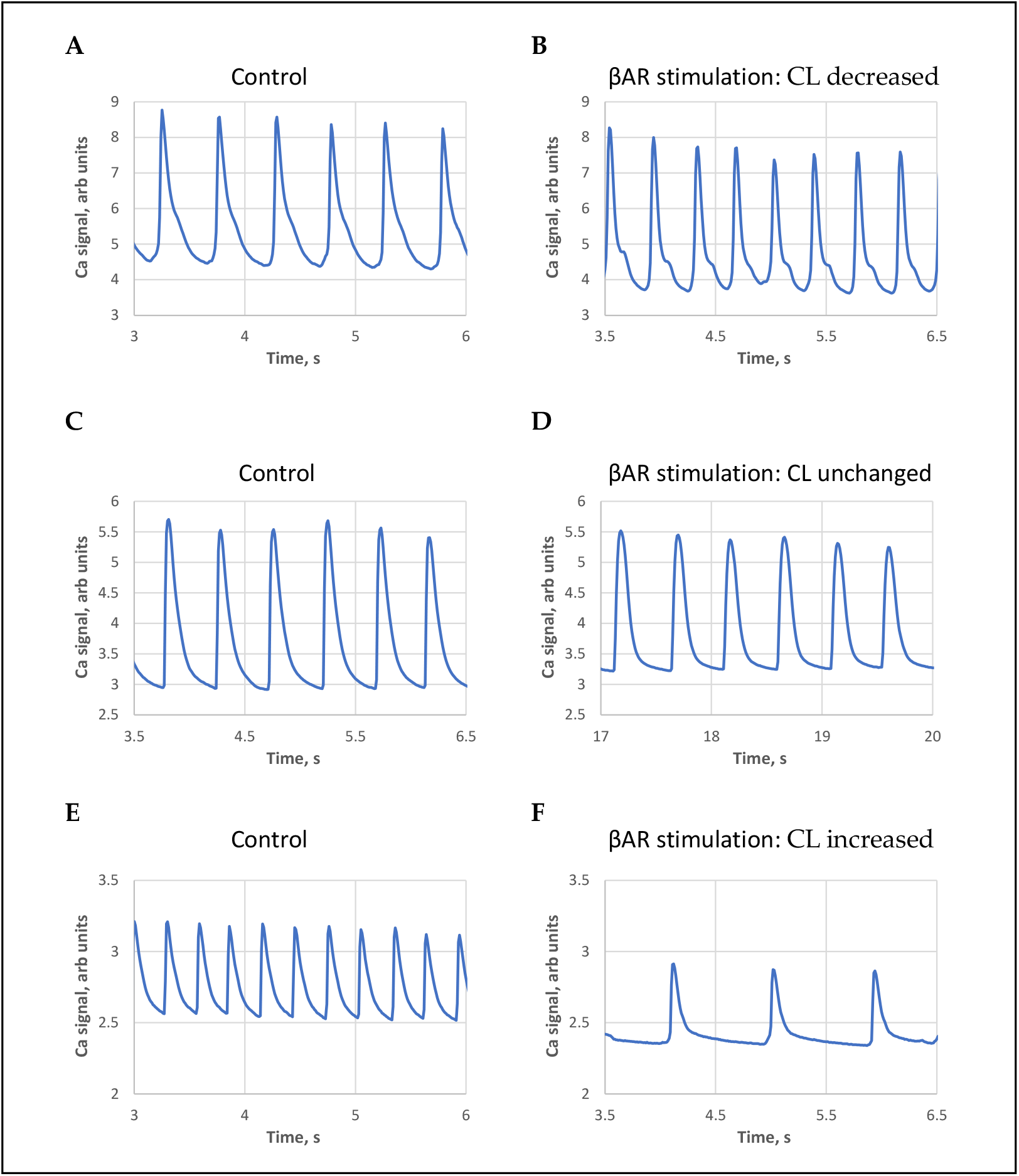
Examples of three types of βAR stimulation effects on AP-induced Ca transient cycle length (CL): (A,B): decrease; (C,D): no change; and (E,F): increase. All traces have the same 3s duration for clear visual comparison. See also respective Videos S1-S6 of cell Ca dynamics in control and in the presence of βAR stimulation.

We noticed that the classical response (positive in terms of AP rate increase) was typically observed in SAN cells with larger basal cycle lengths. Contrastingly, the paradoxical response (cycle length increase) was mainly observed in cells with short baseline cycle lengths, i.e. in cells with higher basal beating rates. The largest cycle length shortenings were observed in cells operating at extremely long cycle lengths (Figure 2, Videos S7 and S8), i.e. in SAN cells bordering on the dormant state [10–12].

**Figure 2.**
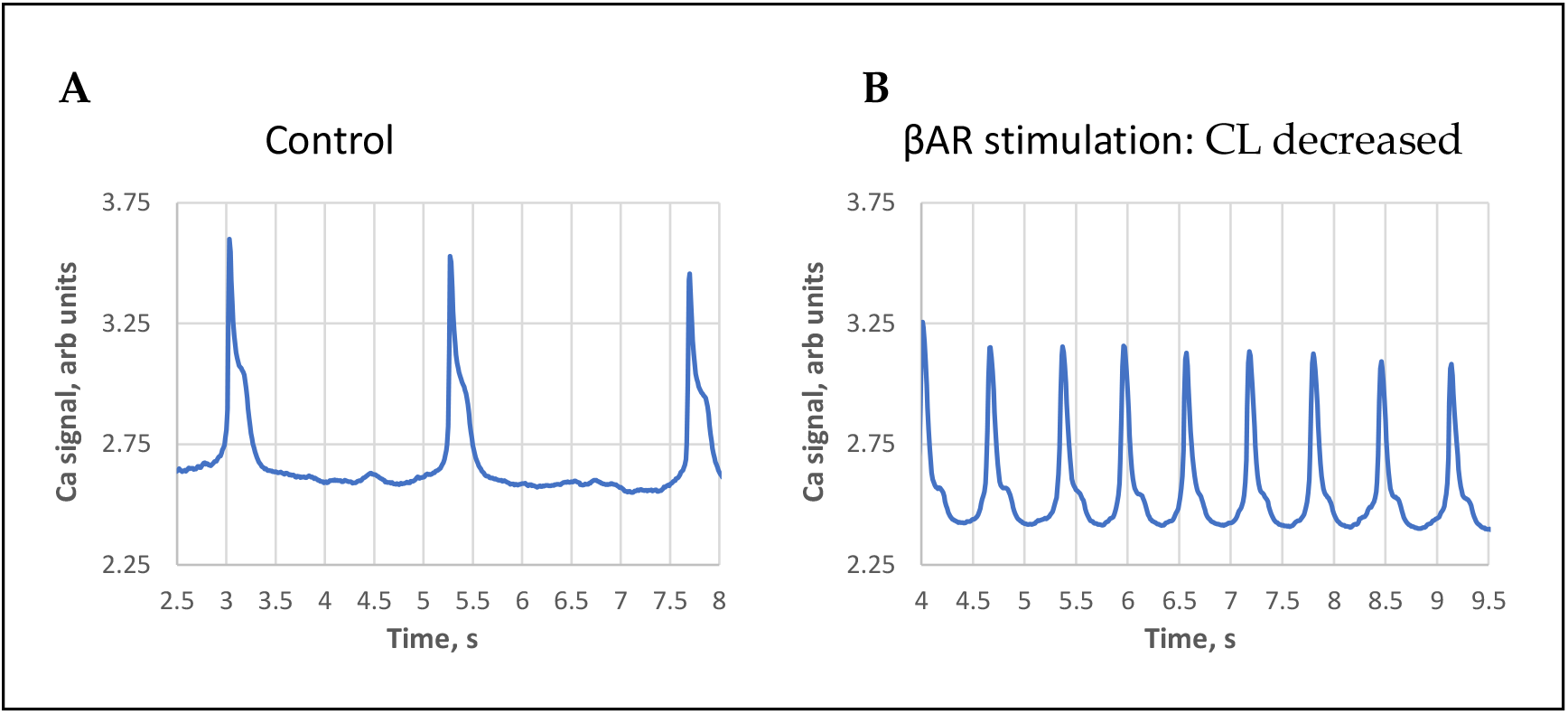
An example of βAR stimulation effect in cells with extremely long AP-induced Ca transient cycle length (CL). (A): at the base line, i.e. “almost dormant cell”. (B): βAR stimulation. Both traces have the same 5.5 s duration for clear visual comparison. See also respective Videos S7-S8 of cell Ca dynamics in control and in the presence of βAR stimulation.

We illustrated and statistically evaluated this result by plotting cycle length change vs. initial cycle length before βAR stimulation for each cell tested (Figure 3). The relationship was closely described by a linear function, with R^2^=0.961. The fitted line crossed the x axis at a cycle length of 491 ms (“no change” cycle length), providing an approximate border between negative and positive responses that is also close to the average cycle length (507 ms) in the presence of βAR stimulation. This suggests synchronization (or grouping) of the responses around this no-change cycle length value. Such synchronization is also evidenced by a substantially smaller standard deviation of cycle lengths in βAR stimulated cells. Synchronization is also suggested by the substantially smaller coefficient of variation (CV) (33% vs. 93%, Table 1) of cycle length in the presence of βAR stimulation.

**Figure 3.**
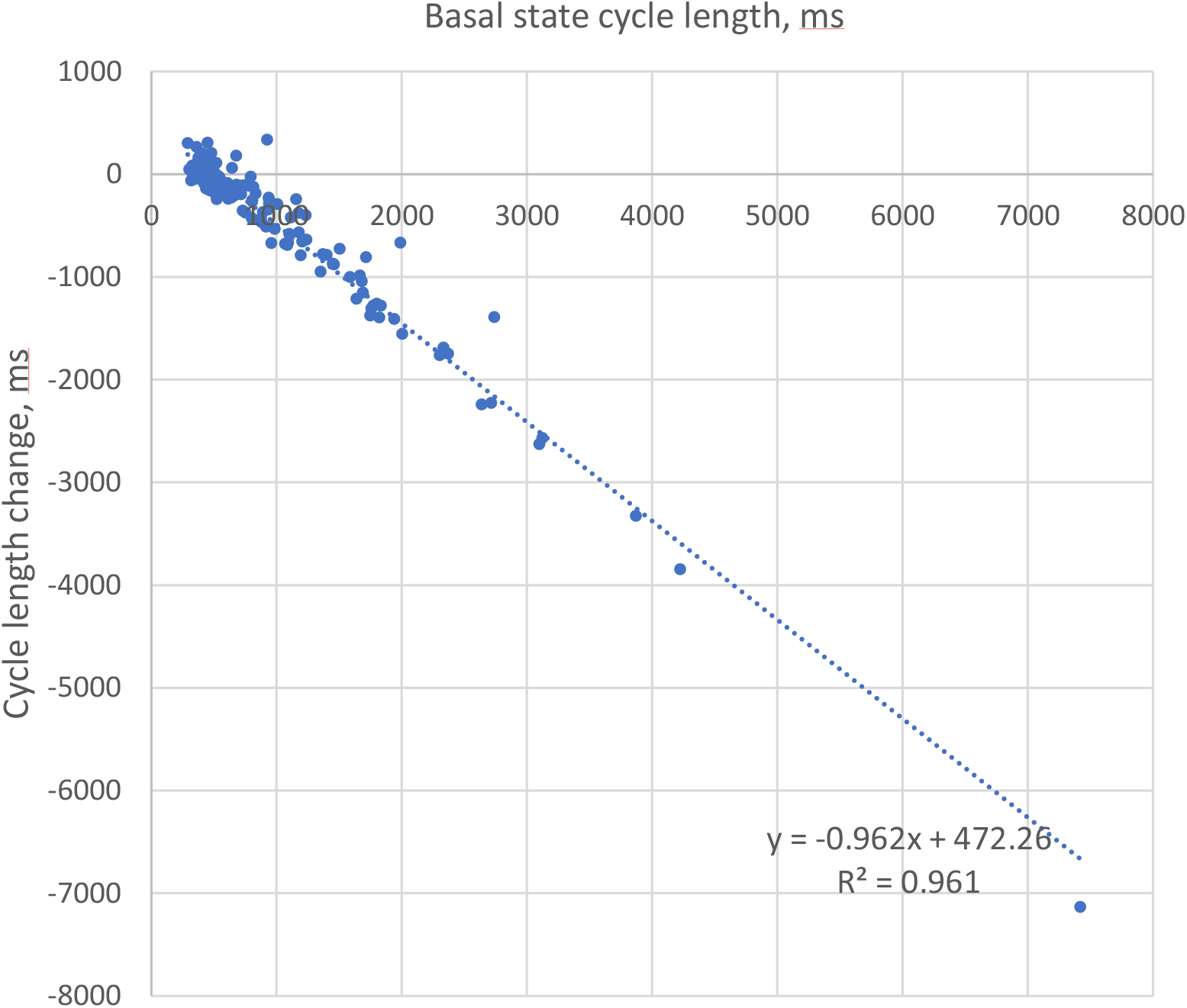
Strong correlation of the initial cycle length (basal state) with change in the cycle length in the presence of βAR stimulation. Negative numbers on Y axis represent cycle length decrease, i.e. rate increase.

The βAR stimulation associated cycle length synchronization is illustrated by comparison of the respective histograms for cycle lengths in control and in the presence of βAR stimulation (Figure 4). Cycle length distribution is spread widely in the basal state, with many cells exhibiting long, some extremely long values. In the presence of βAR stimulation the distribution substantially narrows, and almost all cells operate around a common, higher (on average) rate.

**Figure 4.**
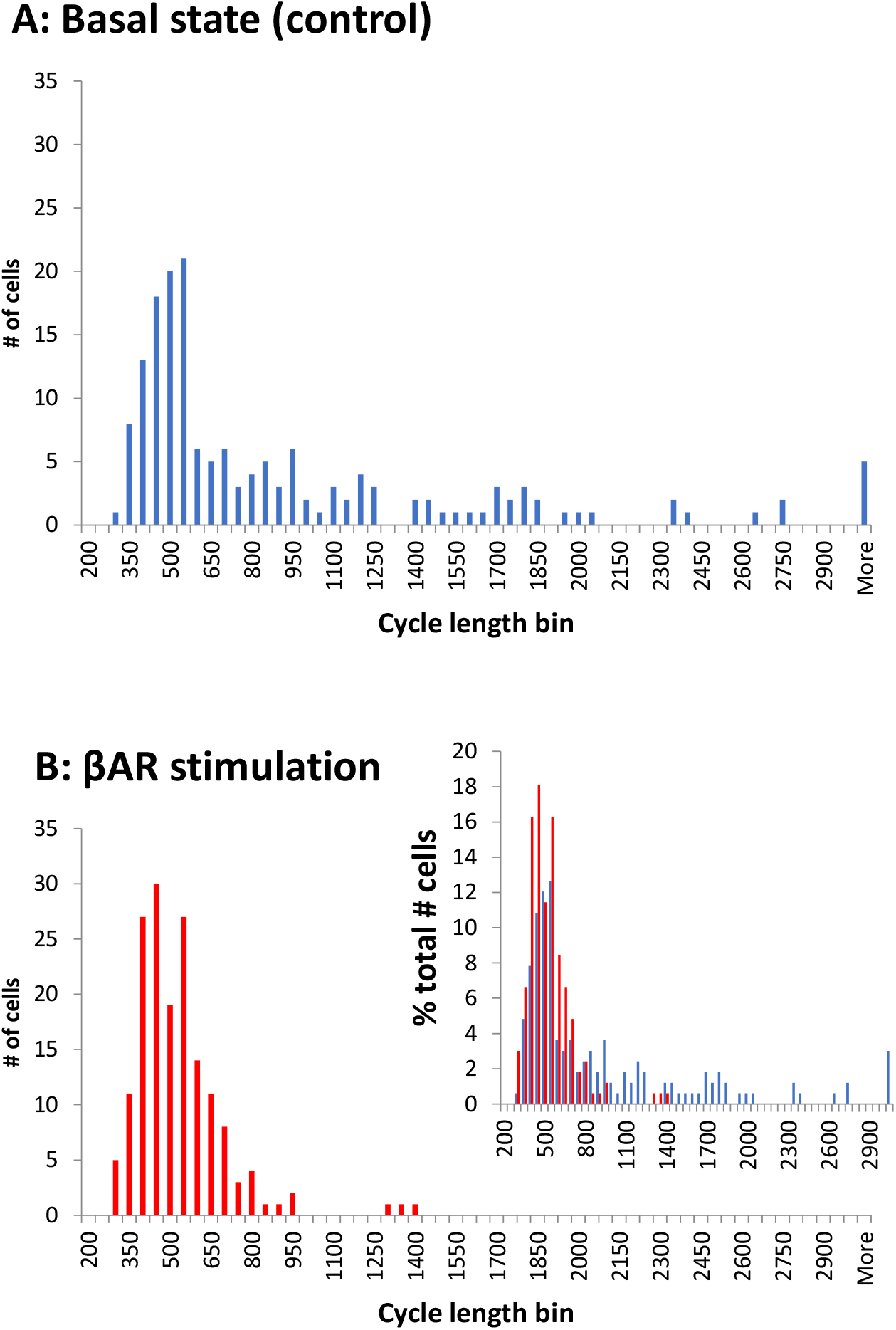
βAR stimulation synchronizes cell rates around a higher (on average) common rate, i.e. shorter cycle length. Histograms show cycle length distributions in 166 cells tested in the present study. Bin size is 50 ms. (A): in control. (B): in the presence of βAR stimulation. Inset: overlapping histograms in panels A and B expressed in terms of the percentage of cells in each cycle length bin illustrate a more narrow, synchronized distribution of cycle lengths in stimulated cells (red) relative to control (blue).

We performed additional data analyses to ensure that our results were robust and independent of number of cells measured from each heart. Our measurements varied from a minimum of 1 to a maximum of 19 cells/ animal (see histogram in Figure S3). We virtually divided the entire cell population approximately by half with substantially different numbers of cells/animal: the subgroup 1 has 82 cells with 1 to 7 (cells/animal), and the subgroup 2 has 84 cells with 8 to 19 (cells/animal). Both subgroups have similar average cycle lengths in basal state and during βAR stimulation, similar fractions of cells with cycle length increase, and similar results of linear recreation analyses vs. each other and vs. all cells measured (Table S1, Figure S4). Thus, the results of our study are robust and independent of cells measured from each isolation.

### 3.2. LCR changes

Our previous studies have shown the crucial importance of LCRs for facilitating the effect of βAR stimulation in AP firing SAN cells [25], and in dormant SAN cells [10–12]. We consequently measured and compared changes in key LCR characteristics in contrasting cells responding to βAR stimulation with cycle length decrease vs. cycle length increase. All key LCR characteristics in SAN cells responding with cycle length decrease changed substantially (Figure 5) in line with the coupled-clock theory (see more in Discussion). In contrast, LCR characteristics in cells exhibiting cycle length increase showed no significant change, except the LCR period, which increased (Figure 6).

**Figure 5.**
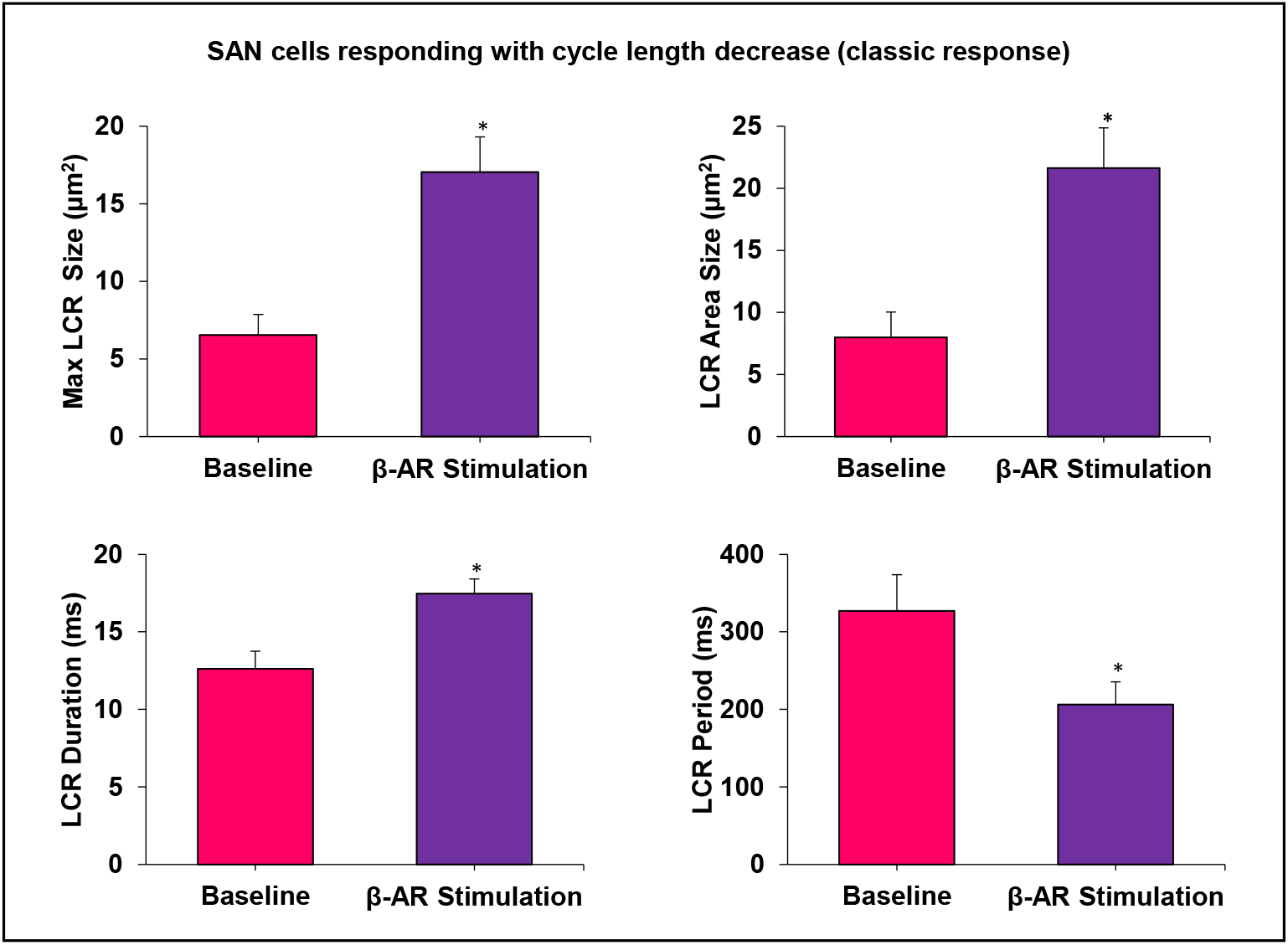
All measured LCR characteristics substantially and significantly change in cells with cycle length decrease (rate increase, i.e. classic response). *P<0.05, n=4 cells.

**Figure 6.**
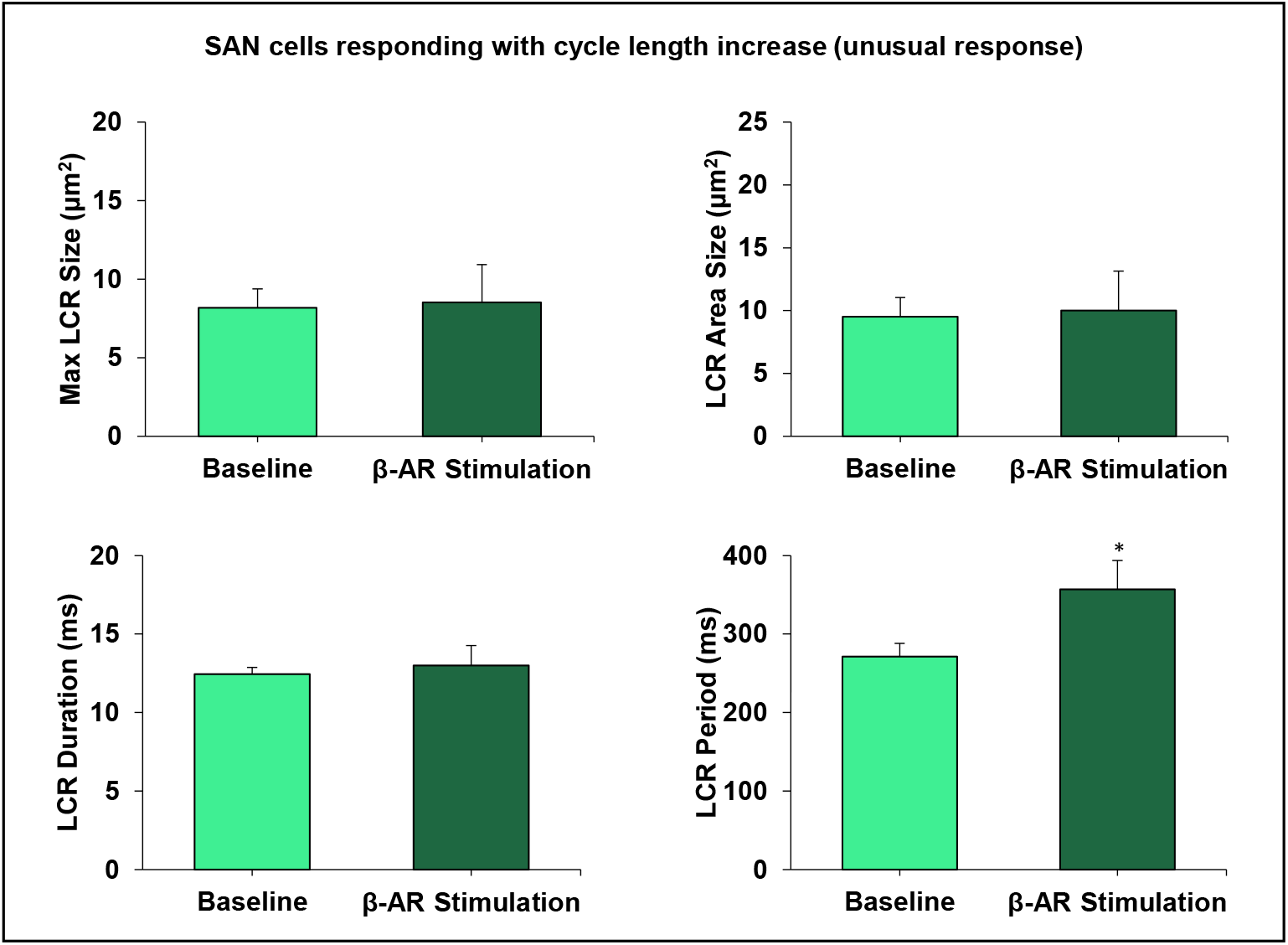
LCR characteristics do not change, except LCR period, in cells responding with the cycle length increase (rate decrease, i.e. a new response type). * P<0.05, n=5 cells.

## 4. Discussion

### 4.1. Result summary

The present study, for the first time, has examined the diverse effects of βAR stimulation in a large number of isolated SAN pacemaker cells (n=166). Previous studies have evaluated the average effect of βAR stimulation in a small population of cells that were spontaneously beating after isolation, with a rate similar to that of the intact SAN. Recent findings in the intact SAN as a whole [8,9], and in dormant isolated cells [10–12] have inspired us to investigate *all* SAN cells isolated from SAN tissue, to better appreciate their behavior and understand their possible contribution to automaticity. Our novel aim here was to examine βAR stimulation effects in a large population of SAN cells generating spontaneous AP-induced Ca transients *at any rate*, including very low rates, bordering on dormancy. We tested the hypothesis that the responses would vary substantially among cells, and endeavored to find a general relationship of the rate change during βAR stimulation among cells. We found that βAR stimulation responses did indeed markedly vary. Furthermore, the cycle length change in each cell was highly correlated (R^2^=0.97) with the cycle length before stimulation. While on average, cycle length decreased, larger cycle length shortenings were observed in cells with longer cycle lengths before stimulation. Vice versa, cells with shorter cycle length showed smaller responses or even paradoxical cycle length increases (Figure 3). Thus, βAR stimulation critically synchronizes the intrinsic operation of SAN cells towards a higher average rate, differing fundamentally from the dogma of a uniform shift in each cell [2].

### 4.2. Possible mechanisms underlying different cell sensitivity to βAR stimulation

The large spectrum of cell responses to βAR stimulation is likely linked to the extremely high heterogeneity of SAN cells with respect to their expression of key proteins (including Ca cycling proteins [17,26]), and ion current densities [17–19]. Indeed, our theoretical studies [27] employing wide ranging, multidimensional sensitivity analyses tested hundreds of thousands of numerical models with different yet realistic parameters of both Ca and membrane clocks. These analyses demonstrated substantial variation in the ranges of βAR stimulation effect. A major conclusion (related to this study) was that the full physiological range of rates can be achieved only in SAN cell models that incorporate a Ca clock. The membrane and Ca clocks operate in synergy providing robust, flexible, and energy efficient spontaneous AP firing [13,27]. In mathematical terms, the Ca clock operates *via* a *criticality* mechanism [28], employing phase-like transitions [29]. It is continuously but variably coupled to the membrane clock that generates current/voltage oscillations *via* a *limit-cycle* mechanism [30]. The criticality mechanisms, in turn, are governed by a power law and self-similarity across wide scales [31]. Nature harnesses these universal and synergetic mechanisms to effectively execute the wide range of autonomic modulation receptor stimulation [32] observed in species with substantially different ‘normal’ heart rates [33].

The specific mechanism of the beating rate increase during βAR stimulation in our study is also likely linked to coupled-clock function. This is evidenced by respective changes in LCR characteristics (Figure 5) including an increase in average LCR area, maximum LCR size and LCR duration. These all indicate a stronger interaction of LCRs with the membrane clock and a larger total interaction area with Na/Ca exchanger. This LCR-linked mechanism is similar to that we reported for βAR stimulation in dormant cells [10–12] (discussed in Introduction).

Some cells (mainly with high basal rate) responded with rate slowing during βAR stimulation (Figure 1E), with their LCR period significantly increasing (Figure 6). An increase in LCR period is a common feature of SAN cells that decrease their firing rate (e.g. LCR period increases in the presence of cholinergic receptor stimulation [34]). Thus, the rate decrease could be LCR-dependent due to shift in the timing of LCR occurrences. Other LCR characteristics in these cells (assessing strength of LCR signal) remained unchanged (Figure 6). This indicates that the Ca clock (and its coupling to the membrane clock) is tuned to operate differently in these cell populations. This tuning can be linked to the specific settings of complex biochemical reactions (the “biochemical engine” of the cardiac pacemaker). These determine, among other things, cAMP balance and protein phosphorylation driving the Ca clock and the coupled-clock system [14]. The biochemical engine can be slowed down *via* appropriate “brakes”. These include the wide spectrum of different adenylyl cyclases (e.g. Ca-inhibited types), phosphatases, and phosphodiesterases, their operating regimes, and interplay including local interactions, e.g. in microdomains [35,36]. Bradycardic mechanisms of the membrane clock that may contribute to the rate slowing include Ca-activated K channels and Ca-dependent inactivation of ICaL.

### 4.3. A broader interpretation of our results towards the entire range of autonomic modulation, including dormant cells

In the broader context, our finding that βAR stimulation synchronizes intrinsic cell oscillations (i.e. spontaneous AP rates of cells in isolation) may be extended to the entire range of SAN beating rates executed by autonomic modulation. This includes cholinergic receptor stimulation, but with the opposite effect, to desynchronize cell function. In the present paper, in the basal state, the distribution of intrinsic rates is dispersed, but still has a notable peak (Figure 4, top panel). Cell synchronization within the cell population in the presence of βAR stimulation transforms the distribution into a single narrow peak (Figure 4, bottom panel). Cell desynchronization in the presence of cholinergic receptor stimulation would likely result in the opposite effect, i.e. a more disperse (disordered) distribution than in the basal state. Indeed, SAN cells isolated from rabbit hearts substantially decrease or increase their AP cycle length standard deviation upon βAR or cholinergic receptor stimulation, respectively [32].

If this hypothetical synchronization/desynchronization mechanism of cell population redistribution is correct, one could expect that low-rate cells would become non-firing cells in the presence of cholinergic stimulation (following Fenske et al. hypothesis), whereas in the presence of βAR stimulation low-rate cells would become higher-rate cells (present study, Figure 4) and non-firing/dormant cells become firing cells [10–12]. Thus, our broader interpretation of autonomic modulation at the level of the SAN cell population includes redistribution of intrinsic cell functions within the entire spectrum of rates, including in cells beating with a rate of zero, i.e. dormant cells. As such, we hypothesize that dormant cells represent an indispensable part of the SAN cell spectrum. This view is fundamentally different from the old-standing dogma of rate regulation *via* “the pacemaker cell”, i.e. a hypothetical, average SAN cell executing the entire range of autonomic modulation [37–41].

### 4.4. A broader interpretation of our results towards synchronization mechanisms of cell signaling in intact SAN: cell specialization and local control by autonomic system

The findings of the present study are in line with the results of a preliminary study[42] that cells isolated from different SAN regions exhibit not only different basal rates, but also different responses to βAR stimulation. Different cell responses to βAR stimulation, in turn, suggests that some SAN cells are specialized to execute increases in heart rate. Indeed, recent studies [43] discovered two competing right atrial pacemakers - one near the superior vena cava and one near the inferior vena cava. These preferentially control fast and slow heart rates. Cholinergic receptor stimulation causes some SAN cells to become silent, which contributes to rate decrease as proposed by Fenske et al. [9]. Still other areas may be unresponsive to acetylcholine [44]. Using small tissue pieces dissected from rabbit SA node, Opthof et al. [45] found that SA node pieces bordering the left atrium were quiescent when dissected, but became activated to fire APs in the presence of adrenaline or acetylcholine.

Cell specialization to generate different rates and different sensitivity of cells to autonomic stimulation within the cellular network of intact SAN can be interpreted to indicate that the role of autonomic modulation is to *redistribute signaling* (i.e. local control) within the cellular network. This is likely to amplify the contribution and deliver oscillations of a particular rate generated by a specialized cell population towards the overall control of SAN output. Such signal amplification within the intact SAN may be executed *via* stochastic resonance [46] and/or percolation phase transition [30]. Another potential amplification mechanism is linked to dynamic changes in the SAN network itself, i.e. autonomic system-guided local control of dynamic signal channeling. Indeed, during βAR stimulation, some additional (multiple) signaling pathways may be created within the network, as more non-firing/dormant cells become firing cells. On the other hand, those cells that initially had faster intrinsic rates but slowed down in response to βAR stimulation (about 23%, Table S1) may impede certain signal transduction (or propagation) pathways, preventing reentries, thus acting as a safety mechanism.

Thus, cells not only synchronize their intrinsic firing rates (present study), but also their synchronized firing is more robustly, synchronously, and safely delivered to SAN output within the tissue *via* dynamic signal channeling. In contrast, some signal paths to SAN output may be blocked by temporary nonfiring/dormant cells during the cholinergic response (in addition to the tonic entrainment mechanism of Fenske at al. [9]). These may become available once more in the basal state firing. Such a specialized cell population-based mechanism (autonomic system local control of signal channeling) is reflected by the classic phenomenon, known as the shift of the dominant leading pacemaker site [47].

Interestingly, respiration arises from complex and dynamic neuronal interactions within a diverse neural microcircuit, and pacemaker properties are not necessary for respiratory rhythmogenesis [48]. Thus, regulation of cardiac automaticity can also theoretically utilize such a neural microcircuit mechanism. How specifically, cell specialization and different degrees of synchronization in intrinsic cell signaling would be guided/modulated by autonomic system (locally and over entire SAN) and translated into a higher or lower rate of impulses emanating from the SAN remains unknown and merits further study. A problem with interpretation of cell population data to the network level at this moment is that the new paradigm of heterogeneous signaling (including subthreshold signals, non-firing cells, and cells operating at various rates) has only recently been proposed [8,9]. While it has been noted that cell signaling in the SAN network resembles multiscale complex processes of impulse generation within clusters of neurons in neuronal networks [8], establishing exact mechanisms of this complex signaling represents **the frontier of cardiac pacemaker research**. Testing new ideas and establishing exact mechanisms require new theoretical modeling. Multicellular numerical models of SAN function have been developed in 1 dimension (as a linear chain of consecutive SAN cells) [49–54], in 2 dimensions [55–57] and even in 3 dimensions [58]. Those models have provided new insights about the importance of cell interactions, but they have not tested importance of dormant cells (with their subthreshold signaling) [10–12], cells operating at low rates found in intact SAN [8], and autonomic modulation. The cell population data presented here will be helpful for future numerical studies of SAN tissue to decrease uncertainties in choosing cell model parameters.

### 4.5. Importance of beat-to-beat variability

While the present study was focused on the rate responses, another important aspect of function of SAN cells is their beat-to-beat variability that ultimately contributes to heart rate variability. In addition to respective rate changes, βAR stimulation increases rhythmicity of AP firing (*via* cell firing synchronization), whereas cholinergic receptor stimulation decreases rhythmicity of spontaneous AP firing [32,59]. Furthermore, Ca and membrane potential transitions during APs are found to be self-similar and to the variability of AP firing intervals across the entire physiologic range during autonomic receptor stimulation [32]. In fact, the rate and rhythm are intrinsically linked to each other at each and all scale levels (single cells, tissue, and heart) [59]. The data from the present study provides a new insight into this phenomenon. If the SAN rate and rhythm result from an integration of signals from many or all cells, the network of cells with a narrower distribution of intrinsic rates in the presence of βAR stimulation (Figure 4), would logically give less variability at the output after the integration. Thus, future experimental studies of diverse populations of SAN cells and respective theoretical approaches will be more enlightening if they include both rate and rhythm considerations.

### 4.6. Functional benefits of heterogeneity at tissue level

SAN cells exhibit extremely heterogeneous morphological, biophysical, and biochemical properties. Why? What are the benefits and importance of this heterogeneity among cells within the SAN tissue? A comprehensive summary of thoughts in this regard was presented about 20 years ago in a seminal review by Boyett et al. [47] that concluded: “The heterogeneity is important for the dependable functioning of the SA node as the pacemaker for the heart, because (i) *via* multiple mechanisms, it allows the SA node to drive the surrounding atrial muscle without being suppressed electrotonically; (ii) *via* an action potential duration gradient and a conduction block zone, it promotes antegrade propagation of excitation from the SA node to the right atrium and prevents reentry of excitation; and (iii) *via* pacemaker shift, it allows pacemaking to continue under diverse pathophysiological circumstances.” The functional importance of heterogeneity in cellular networks has been also demonstrated in theoretical studies. For example, the presence of the Ca clock in addition to the membrane clock protects the SAN from annihilation, associated with sinus node arrest [54]. The network structure critically determines oscillation regularity [60] and random parameter heterogeneity among oscillators can consistently rescue the system from losing synchrony [61].

Another possible advantage of cell specialization and heterogeneous signaling among SAN cells is more efficient energy consumption. For example, cells specializing in impulse conduction to the atria, building a “conduction block zone”, or generating a particular rate, may not possess a “powerful” Ca clock, consuming a lot of energy for its cAMP/PKA signaling [62], required for wide range rate regulation [27]. Furthermore, heterogeneous signaling in SAN tissue [8] implies that during rest periods, many cells may stay in reserve, firing APs at lower rates (and not necessarily rhythmically), or even fall to dormancy, consuming much less energy. For example, in cats, the primary pacemaker consists of at most 2000 cells, but appears to function normally with less than 500 cells [63]. During the fight-or-flight response, however, we would expect that all reserves are recruited and maximally synchronized to generate rhythmic frequent impulses for maximum blood supply for the sake of survival at any energy cost.

Yet another level of heterogeneity and complexity of cardiac pacemaker tissue is its intense neuronal innervation network [64–66], dubbed the “little brain of the heart” [67]. This provides precise control of SAN pacemaker cell function (*via* possible local control and guidance of heterogeneous signaling discussed above). SAN pacemaker cells also have interactions with non-excitable cells, such as telocytes [68,69] and fibroblasts [70] that could also aid in local control of heterogeneous signaling. Fibroblasts provide obstacles, ion current sinks or shunts to impulse conduction depending on their orientation, density, and coupling [51].

## 5. Conclusions

The response of isolated spontaneously beating SAN cells to βAR stimulation is diverse, and besides cells which increase their beating rates, includes cells that are unaffected, and some which paradoxically slow down. Individual cell response depends on the basal rate of beating, and overall there is a synchronization in beating rates in response to βAR stimulation across the cell population. The effect is mitigated through the coupled clock system, but effects on LCRs differ in cells depending on their response to βAR stimulation. Understanding how markedly heterogenous populations of SAN cells respond to autonomic stimulation, and how this is integrated into the function and output of the SAN at the tissue and organ level represents one critical frontier in SAN research mandating intense future study.

## Supporting information

Video S1 for Fig1A, control

Video S2 for Fig1B, isoproterenol

Video S3 for Fig1C, control

Video S4 for Fig1D, isoproterenol

Video S5 for Fig1E, control

Video S6 for Fig1F, isoproterenol

Video 7 for Fig2A, control

Video 8 for Fig2B, isoproterenol

## Supplementary Materials

**Supplemental Figures S1-S4**

**Supplemental Table S1**

**Supplemental Videos:**

**Video S1:** An example of classical type of βAR stimulation effects to decrease cycle length (i.e. to increase cycling rate). This 3.67 s video shows a 2D recording of Ca signals in basal state before stimulation. See Figure 1A.

**Video S2:** An example of classical type of βAR stimulation effects to decrease cycle length (i.e. to increase cycling rate). This 3.67 s video shows a 2D recording of Ca signals of the same SA node cell as in Video S1 in the presence of stimulation. See Figure 1B.

**Video S3:** An example of absence of notable βAR stimulation effect on cycle length. This 3.5 s video shows a 2D recording of Ca signals in basal state before stimulation. See Figure 1C.

**Video S4:** An example of absence of notable βAR stimulation effect on cycle length. This 3.5 s video shows a 2D recording of Ca signals in the presence of stimulation. See Figure 1D.

**Video S5:** An example of unusual βAR stimulation effect to increase cycle length (i.e. decrease cycling rate). This 3.67 s video shows a 2D recording of Ca signals in basal state before stimulation. See Figure 1E.

**Video S6:** An example of unusual βAR stimulation effect to increase cycle length (i.e. decrease cycling rate). This 3.67 s video shows a 2D recording of Ca signals in the presence of stimulation. See Figure 1F.

**Video S7:** An example of βAR stimulation effect to substantially decrease cycle length in a cell with extremely long cycle length in the basal state. This 5.96 s video shows a 2D recording of Ca signals in basal state before stimulation. See Figure 2A.

**Video S8:** An example of βAR stimulation effect to substantially decrease cycle length in a cell with extremely long cycle length in the basal state. This 5.96 s video shows a 2D recording of Ca signals in the presence of stimulation. See Figure 2B.

## Author Contributions

Conceptualization, VAM and EGL; methodology, MSK, OM, VAM; software, VAM; validation, OM, VAM, EGL; investigation, MSK, LAM, OM, VAM; resources, EGL; data curation, VAM; formal analysis, LAM, MSK; writing—original draft preparation, MSK, VAM; writing—review and editing, MSK, OM, LAM, EGL, VAM; visualization, VAM, LAM; supervision, EGL, VAM; project administration, EGL. All authors have read and agreed to the published version of the manuscript.”

## Institutional Review Board Statement

The study was conducted in accordance with NIH guidelines for the care and use of animals, protocol # 034-LCS-2019.

## Informed Consent Statement

Not applicable.

## Data Availability Statement

All data are available upon the request sent to the corresponding author (VAM).

## Acknowledgments

The authors wish to acknowledge the assistance of Brice D. Ziman for skillful isolation of SAN cells.

## Abbreviations

SAN: sinoatrial node
AP: action potential
LCR: local Ca release
βAR: β-adrenergic receptor
HCN4: Hyperpolarization Activated Cyclic Nucleotide Gated Potassium Channel 4
I_f_: the “Funny” current, a mixed Na-K inward HCN current activated by hyperpolarization
I_CaL_: L-type Ca current
I_K_: delayed rectifier K current
RyR2: Ryanodine Receptor type 2
SERCA2: sarco/endoplasmic reticulum Ca-ATPase type 2
Cycle length: action potential cycle length
cAMP: cyclic adenosine monophosphate
PKA: cAMP-dependent protein kinase type A

