## Supplemental Figures S1-S4 for "β-Adrenergic Stimulation Synchronizes a Broad Spectrum of Action Potential Firing Rates of Cardiac Pacemaker Cells towards a Higher Population Average"

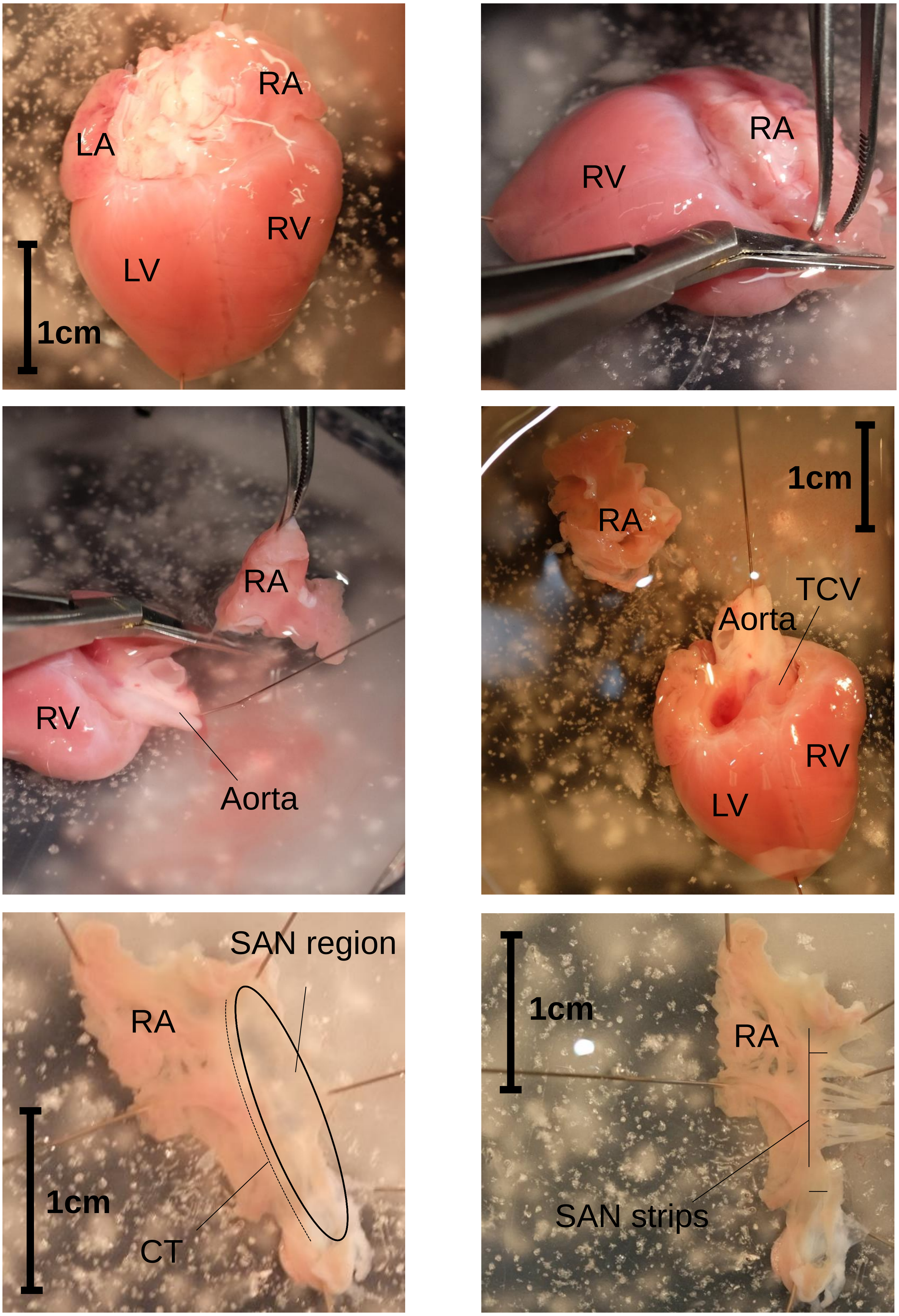


Figure S1: Location of guinea pig SA node in relation to the whole heart during our isolation procedure. LA=left atrium; LV=left ventricle; RA=right atrium; RV=right ventricle; TCV=tricuspid valve; CT=cristae terminalis; SAN=sinoatrial node. Modified from Kim et al. Cell calcium 2018, 74, 168-179, doi:10.1016/j.ceca.2018.07.002

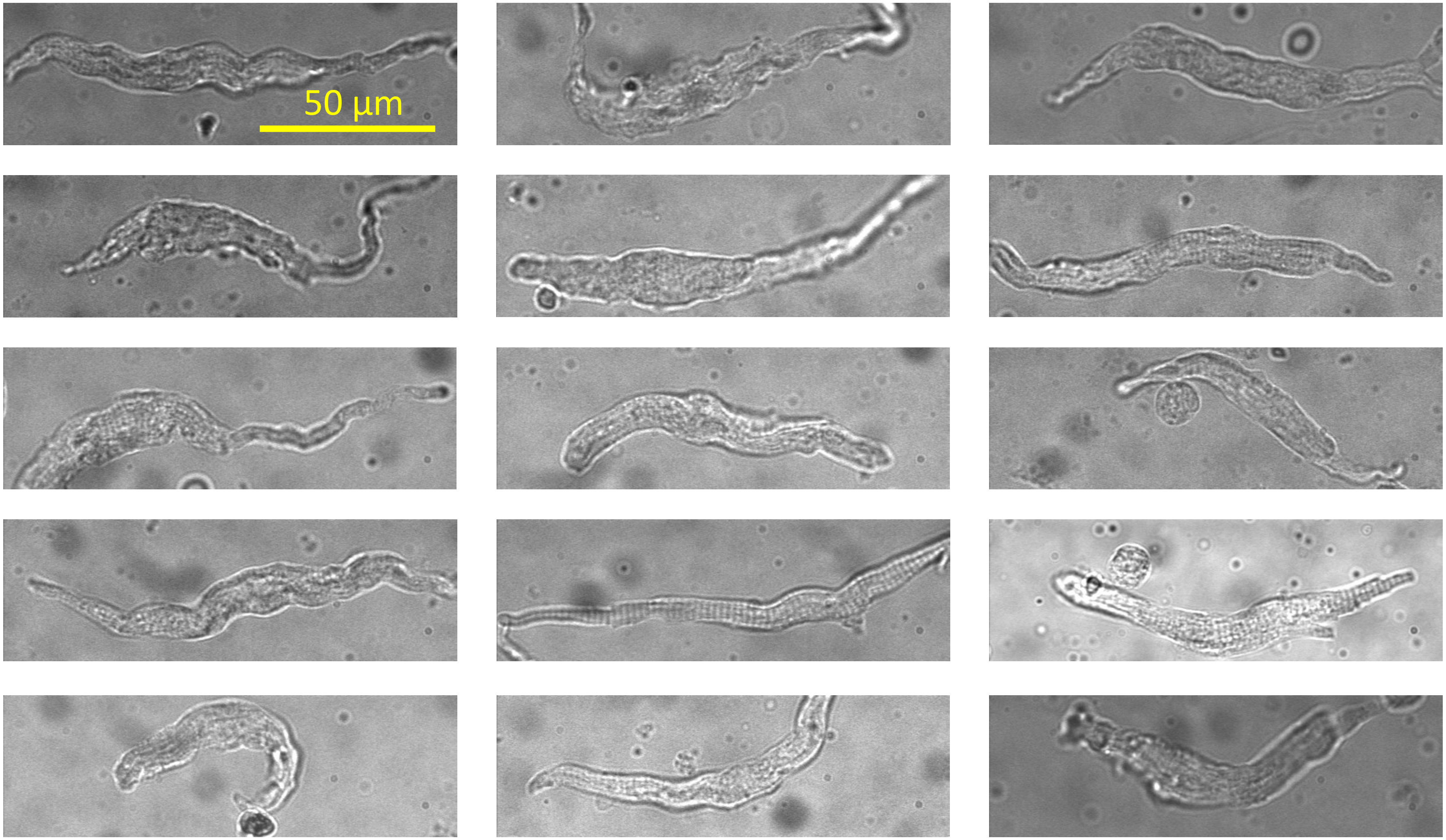


Figure S2: Typical morphologies of cells used in the present study.

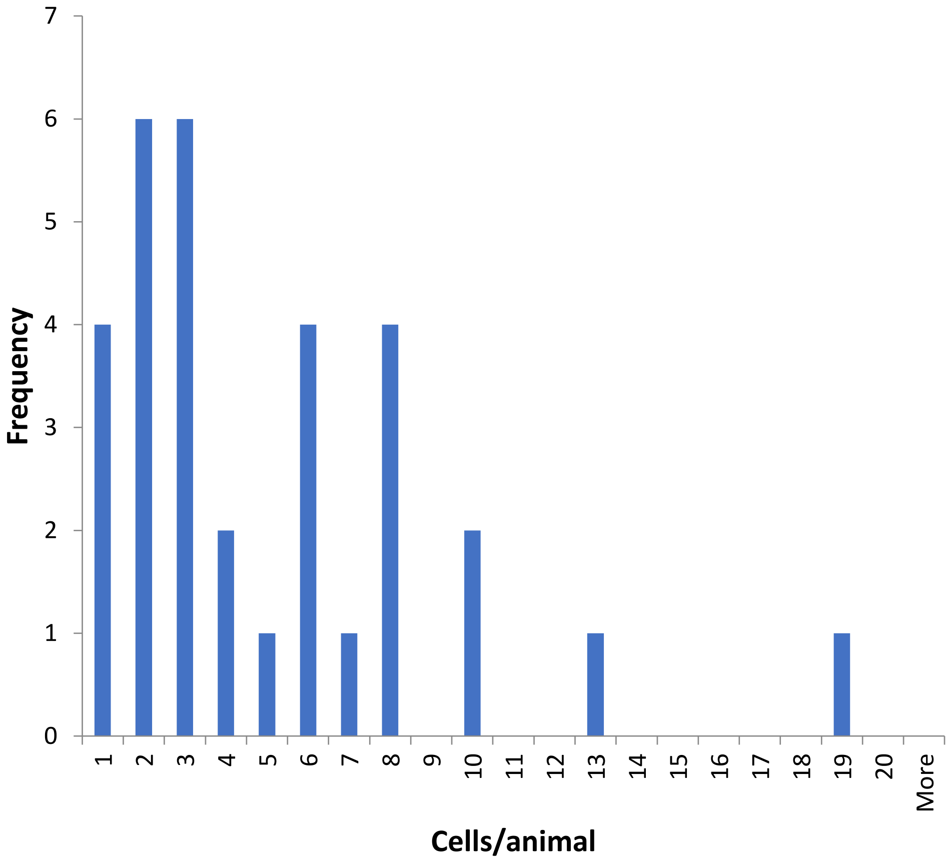


Figure S3: Distribution of our measurements in terms of cells/animal

A

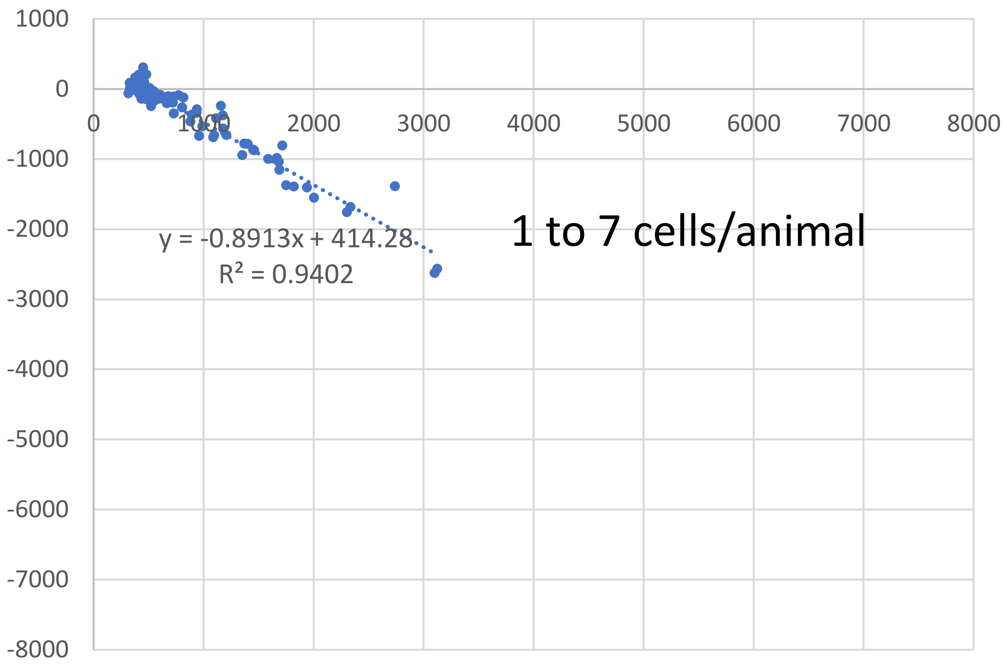


B

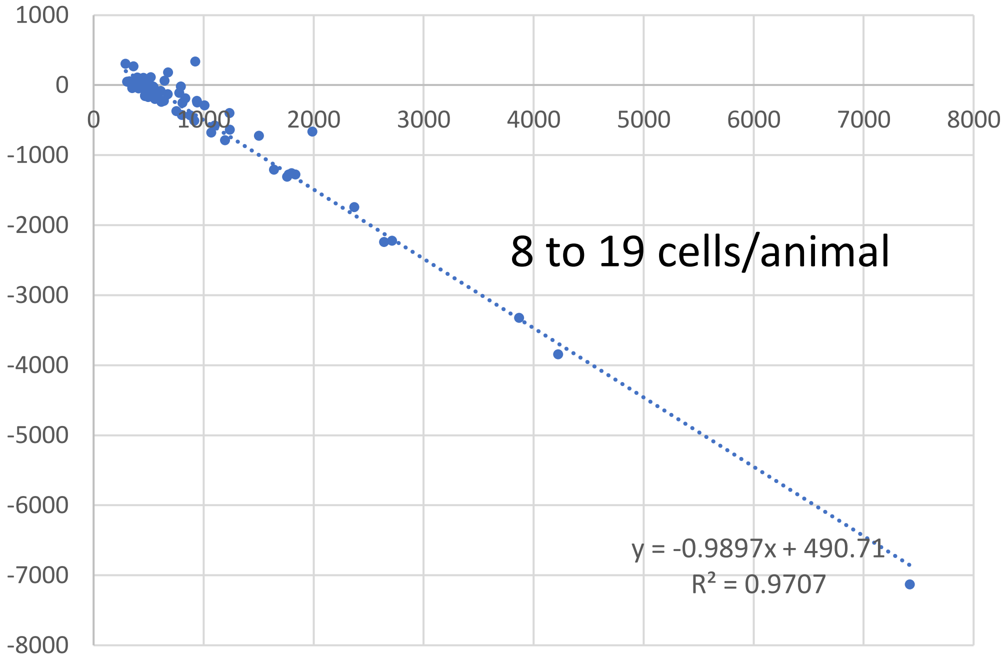


Figure S4: Results of linear regression analyses for subgroup of cells with different number cells measured per animal, illustrating robustness of our findings. (A): Analysis of 82 cells in the subgroup of 1 to 7 cells/animal. (B): Analysis of 84 cells in the subgroup of 8 to19 cells/animal. Regression line (dotted lines) equations are shown together with respective $R^2$ values.
