## Supplemental Table S1 for "β-Adrenergic Stimulation Synchronizes a Broad Spectrum of Action Potential Firing Rates of Cardiac Pacemaker Cells towards a Higher Population Average"

| Parameter | All cells | | Cells/animal <=7 | | Cells/animal >=8 | |
|---|---|---|---|---|---|---|
| Number of cells | 166 | | 82 | | 84 | |
| Conditions | Basal | ISO | Basal | ISO | Basal | ISO |
| Mean value | 921.7614 | 507.3039 | 923.1902 | 514.636 | 920.3667 | 500.1464 |
| SEM | 66.47293 | 13.12734 | 71.66966 | 17.89898 | 111.6491 | 19.24943 |
| SD | 856.4438 | 169.134 | 648.9964 | 162.0821 | 1023.281 | 176.4239 |
| CV% | 92.91382 | 33.33977 | 70.29931 | 31.49452 | 111.1819 | 35.27445 |
| Cells with unusual response (CL increase) | 38 (22.9%) | | 17 (20.7%) | | 21(25%) | |
| Regression slope | -0.962 | | -0.8913 | | 0.9897 | |
| Regression offset | 472.26 | | 414.28 | | 490.71 | |
| $R^2$ | 0.961 | | 0.9402 | | 0.9707 | |

**Table S1.** Statistical data and results of linear regression analyses of AP cycle lengths (given in ms) measured in two subgroups of cells with different numbers of cells/animal compared to the results obtained in all cells. CL, cycle length of AP-induced Ca transient; ISO, isoproterenol; SD, standard deviation; SEM, standard error of mean; CV coefficient of variation (in %) = 100*Mean/SD.
